# Brief report: Attentional Bias towards Social Interactions during Viewing of Naturalistic Scenes

**DOI:** 10.1101/2021.02.26.433078

**Authors:** Simona Skripkauskaite, Ioana Mihai, Kami Koldewyn

## Abstract

Human visual attention is readily captured by the social information in scenes. Multiple studies have shown that social areas of interest (AOIs) such as faces and bodies attract more attention than non-social AOIs (e.g. objects or background). However, whether this attentional bias is moderated by the presence (or absence) of a social interaction remains unclear. Here, the gaze of 70 young adults was tracked during the free viewing of 60 naturalistic scenes. All photographs depicted two people, who were either interacting or not. Analyses of dwell time revealed that more attention was spent on human than background AOIs in the interactive pictures. In non-interactive pictures, however, dwell time did not differ between AOI type. In the time-to-first-fixation analysis, humans always captured attention before other elements of the scene, although this difference was slightly larger in interactive than non-interactive scenes. These findings confirm the existence of a bias towards social information in attentional capture, and suggest that the presence of social interaction may be important in inducing a similar social bias in attentional engagement. Together with previous research using less naturalistic stimuli, these findings suggest that social interactions carry additional social value that guides one’s perceptual system.

## Attentional Bias towards Social Interactions during Viewing of Naturalistic Scenes

Detecting, identifying, and interpreting social cues are important to social success. Research to date shows that our attention prioritizes social information above and beyond low-level physical saliency (e.g. End & Gamer, 2017; Yarbus, 1967). A preference towards attending to other people allows us to quickly and accurately parse social scenes and facilitates social understanding. The interactions between people are a unique source of social cues. Not only do they provide rich and complex information about individuals, but also about the relationships *between* people (Quadflieg & Penton-Voak, 2017). Indeed, from an early age, humans use viewing social exchanges as an opportunity to learn from and about others, including assessing competence (Lee & Rutherford, 2018) and judging the likelihood of affiliation (e.g. Powell & Spelke, 2018). Watching others interacting may also provide a natural opportunity to learn social rules and norms (see Quadflieg & Koldewyn, 2017). Despite the key role interactions play in social understanding, it remains unclear whether that importance is reflected in the degree to which they capture and hold attention above and beyond the simple presence of multiple agents.

Recent evidence suggests that interacting dyads may be perceived and processed differently from either individuals or non-interacting dyads. For instance, pairs of silhouettes that face each other (i.e., interacting) are found and processed more efficiently than either dyads facing away from one another or single individuals in a visual search task (Papeo, Goupil, & Soto-Faraco, 2019). Facing dyads are also recognised as human more quickly, are processed faster and remembered better, and are more susceptible to the inversion effect than non-facing dyads (Papeo & Abassi, 2019; Papeo, Stein, & Soto-Faraco, 2017, Vestner, Tipper, Harlety, Over, & Rueschemeyer, 2019). Looking time data suggests that infants are capable of differentiating between facing and non-facing dyads already at six-months (Papeo, Nicolas, & Hochmann, 2020) and human silhouettes facing one another also draw more visual attention in both neurotypical and autistic children (Stagg, Linnell, & Heaton, 2014). Taken together, these findings suggest that attentional processing within the broader ‘social’ category is altered by the presence of a social interaction. However, it is not yet clear if this effect is purely social, or in part reflects a configurational benefit not specific to social stimuli (e.g., Vestner, Gray, & Cook, 2020).

Importantly, these experiments used tightly controlled stimuli, with figures isolated from both background elements and context. Similarly, they focused on only one (important) cue to social interaction, that of facing direction. In daily life, interactions between people are encountered in any number of contexts, settings, and situations where social information competes for attention with many objects and other distractors. Indeed, one purpose of the attentional bias towards others may, at least in part, be to facilitate our ability to filter out ‘non-social’ elements in order to successfully allocate attention to relevant social information (see Kingstone, 2009; Risko, Laidlaw, Freeth, Foulsham, & Kingstone, 2012). Thus, it is crucial to investigate attention to social interaction within the kinds of complex scenes that are typically encountered in ‘real-life’.

To date, few studies have explicitly investigated the influence of social exchanges on how visual attention operates in complex scenes. Birmingham et al. (2008) looked at visual attentional engagement in scenes depicting either one or three individuals being either active or inactive. Three-person ‘active’ scenes could be either interactive or not. They did not find increased attention towards humans in interactive scenes compared to active but noninteractive scenes, though that particular analysis is only briefly described. In contrast, Villani et al. (2015) found that more attention was paid to the faces and arms of humans in interactive rather than non-interactive paintings, although they did not consider attention to other scene elements. Given these conflicting findings, it remains unclear whether the presence of a social interaction alters attention in complex social scenes.

The current study, thus, aimed to investigate whether the bias to attend to the social elements in naturalistic scenes is moderated by the presence of a social interaction. To address this question, we assessed looking behaviour while participants freely viewed naturalistic photographs depicting two people who were either interacting with each other, or not. Based on the overwhelming previous evidence for a social attentional bias, we expected that humans in the scenes would receive more attentional engagement and faster attentional capture than other elements in the scene (Hypothesis 1). We also hypothesized a stronger social bias via longer engagement with and faster capture towards humans in scenes judged as depicting two interacting than two non-interacting individuals (Hypothesis 2). These data should give us a clear insight into whether the third-person observation of interacting individuals carries important social information over and above the observation of a noninteracting pair.

## Methods

### Participants

An a priori power analysis determined that a sample size of 70 (β=0.80, α=.05) would be sufficient to detect a large effect size (Cohen’s f=.40) in the three-way interaction (G*Power 3.1; Faul, Erdfelder, Buchner, & Lang, 2009). This was pre-registered on AsPredicted (https://aspredicted.org/blind.php?x=se7k7b). In total, 73 participants were recruited for the study through an opportunity sample. Data from two participants who were outside our target age-range, and one who was falling asleep were removed. The final sample of 70 participants (*M*=21.07, *SD*=2.63; 47 females and 1 other) had normal or corrected-to-normal vision. All participants gave informed consent and were reimbursed for their time in the study with course credit or a small monetary reward. The study was approved by the School of Psychology’s ethics board and was conducted in accordance with the 1964 Helsinki declaration.

### Stimuli and Apparatus

Stimuli were presented in a paradigm coded in PsychoPy 2 (Pierce et al, 2019), while data was collecting with an Eyelink Portable Duo eye-tracker (SR Research Ltd., 2012). Stimuli were presented on a 380×215 mm (1920×1080 px) monitor with a grey background. Participants’ binocular gaze^2^ was tracked remotely at a sampling rate of 1000 Hz. Target stimuli, selected from the SUN database (Xiao, Hays, Ehinger, Oliva, & Torralba, 2010), included 60 photographs depicting two people in naturalistic scenes. They did not include any photographs with direct gaze at the camera, interactions with unseen (off-camera) agents, or obvious depictions of emotions. Initially, 127 pictures were chosen and then rated for their interactiveness (1=‘Not at all interactive’, 7=‘Very interactive’) by 26 independent judges. Based on these ratings, the 30 photographs that received the lowest and the 30 that received the highest scores were selected to represent ‘non-interactive’ and ‘interactive’ categories, respectively. All images were pre-processed in Photoshop graphic software (version CC 2019) by removing colour cast and matching the colour scheme to a one of the images (using the ‘match colour’ function) and at a standard size of 860×860 px (13.6×13.6°).

### Procedure

The full experiment consisted of 142 trials in total: 60 trials specific for this experiment and 82 unrelated images for another paradigm. Data from other images will not be presented here. Each participant received on-screen and verbal instructions to simply look at the photographs presented on the screen. Every trial first started with a drift correction dot at the centre of the screen, which remained until participants pressed the ‘space’ button whilst successfully focusing on the dot. The image for that trial was then presented on the left or right side of the screen for 5000ms. The left-most edge of right-hand pictures, and the rightmost edge of left-hand pictures was placed 1° from centre. Whether a picture appeared on the right or the left was counterbalanced between participants and the order of trials was fully randomised for each participant.

### Data Analysis

For each photograph, social and non-social areas of interest (AOI) were defined using the ‘freehand’ function in Eyelink Data Viewer (SR Research Ltd., 2013). The former encompassed all visible body parts (including faces/heads), whereas the latter included everything else in the scene, including background elements (c.f. Fletcher-Watson, Leekam, Benson, Frank, & Findlay, 2009). Dwell time (the amount of time, including fixations and saccades, spent gazing inside that particular AOI) was extracted for both social and non-social AOIs as a measure of general attentional engagement with that AOI. Time-to-first-fixation to social and non-social AOIs was extracted as a measure of attentional capture.

In line with best practice (c.f. Fletcher-Watson et al., 2009), all trials where less than 33% of viewing time was engaged with the target photograph were treated as missing. This included both time when participants might be looking off-screen and times when data was missing (e.g., due to blinks or poor signal). This did not, however, result in the loss of very much data. This procedure resulted in the average exclusion of 0.2% (*M*=0.14, *SD*=0.39, range: 0-2) of trials per participant.

Two different models were then used to assess general attentional engagement and attentional capture, respectively. Dwell time and time-to-first-fixation were analysed using linear mixed-effect (multilevel) modelling with a 2×2 design (nlme package; Pinheiro, Bates, DebRoy, & Sarkar, 2016). Participant information was modelled at the fourth level of the multilevel analysis. Nested within each participant, trial information with the type of scene (interactive or non-interactive) as predictor was modelled at the third level. AOI information with AOI type (social or non-social) as predictor was modelled at the second level. Finally, dwell time or time-to-first-fixation per each AOI was modelled at the first level, nested within each trial. Post-hoc Tukey’s HSD pairwise comparisons were carried out when applicable (emmeans package; Lenth, 2018).

## Results

### Attentional Engagement

Dwell time was examined to determine if the amount of attention paid to social information, in contrast to the rest of the image, differed based on whether the scene was interactive or not. Results showed that the main effect of scene type did not reach significance, *F*(1,69)=0.42,*p=*.517, η^2^p=0.01, confirming that looking time to the picture overall did not differ based on whether it was interactive (*M=*1804.73, *SD*=920.09) or noninteractive (*M*=1782.92, *SD*=928.70). There was, however, a significant main effect of AOI type, *F*(1,138)=18.84,*p<.001*, η^2^p=0.12. Participants on average looked at social information (*M*=1866.17, *SD*=932.92) more than non-social information (*M*=1721.49, *SD*=910.20) across scenes. This effect was moderated by the type of scene (Figure 1), *F*(1,138)=43.37, *p<*.001, η^2^p=.24. Specifically, participants looked at social (*M*=1986.53, *SD*=910.75) more than nonsocial (*M*=1622.93, *SD*=893.27) AOIs in the interactive scenes, *t*(138)=7.73, *p<*.001, *d*=0.66. Yet, in the non-interactive scenes they looked just as long at both social (*M*=1745.58, *SD*=939.49) and non-social AOIs (*M*=1820.25, *SD*=916.49), *t*(138)=−1.59, *p=*.115, *d*=0.14. Indeed, participants numerically looked at non-social elements for longer than social elements in non-interactive scenes, though this difference is too small for us to consider it interpretable.

**Figure 1.**
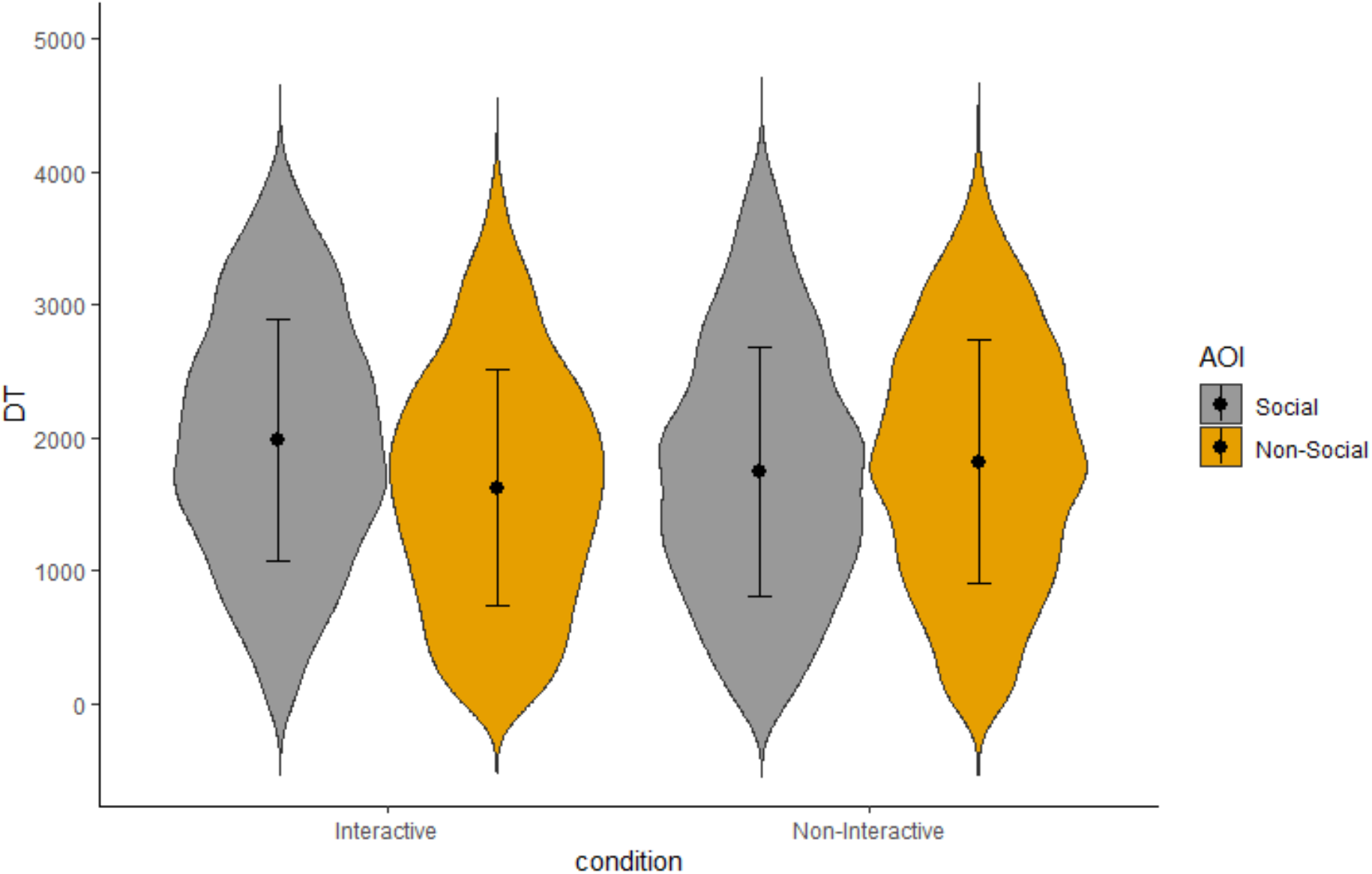
Violin plot for participant’s mean dwell time per AOI and scene type. Error bars represent standard deviations.

### Attentional Capture

Participants’ time-to-first-fixation on social and non-social AOIs were assessed to evaluate potential differences in attentional capture between interactive and non-interactive scenes. Due to never looking at one of the two AOIs, participants were on average missing attention capture data on 1% (*M*=0.30, *SD*=0.55) of social AOIs and 2.77% (*M*=0.83, *SD*=1.14) of non-social AOIs for interactive pictures, *t*(69)=3.45, *p*=.001. For non-interactive pictures, they were missing 2.37% (*M*=0.71, *SD*=0.92) of data for social AOIs and 2.43% (*M*=0.73, *SD*=1.02) for non-social AOIs, *t*(69)=0.09,*p*=.927. For trials with data missing on one AOI, only the data for the available AOI were modelled.

Unsurprisingly, multilevel analysis showed that time-to-first-fixation to the scene overall did not differ based on whether scenes were interactive (*M*=673.33, *SD*=713.73) or non-interactive (*M*=676.40, *SD*=724.94), *F*(1,69)=0.02, *p*=.890, η^2^p<.01. Similar to dwell time analyses, there was both a significant main effect of AOI type [(social: *M*=483.98, *SD*=535.80; non-social: *M*=867.52, *SD*=822.22), *F*(1,138)=423.29, *p*<.001, η^2^p=75] and an interaction effect between AOI and scene type (Figure 2), *F*(1,138)=21.56, *p*<.001, η^2^p=.14. In interactive scenes, participants looked at social AOIs (*M*=440.54, *SD*=462.51) faster than at non-social AOIs (*M*=910.34, *SD*=836.14), *t*(138)=−17.85, *p*<.001, *d*=−1.52. To a lesser extent, their attention was also captured more quickly by social (*M*=528.11, *SD*=598.08) than non-social (*M*=824.77, *SD*=806.03) AOIs in non-interactive scenes, *t*(138)=−11.24, *p*<.001, *d*=−0.97.

**Figure 2.**
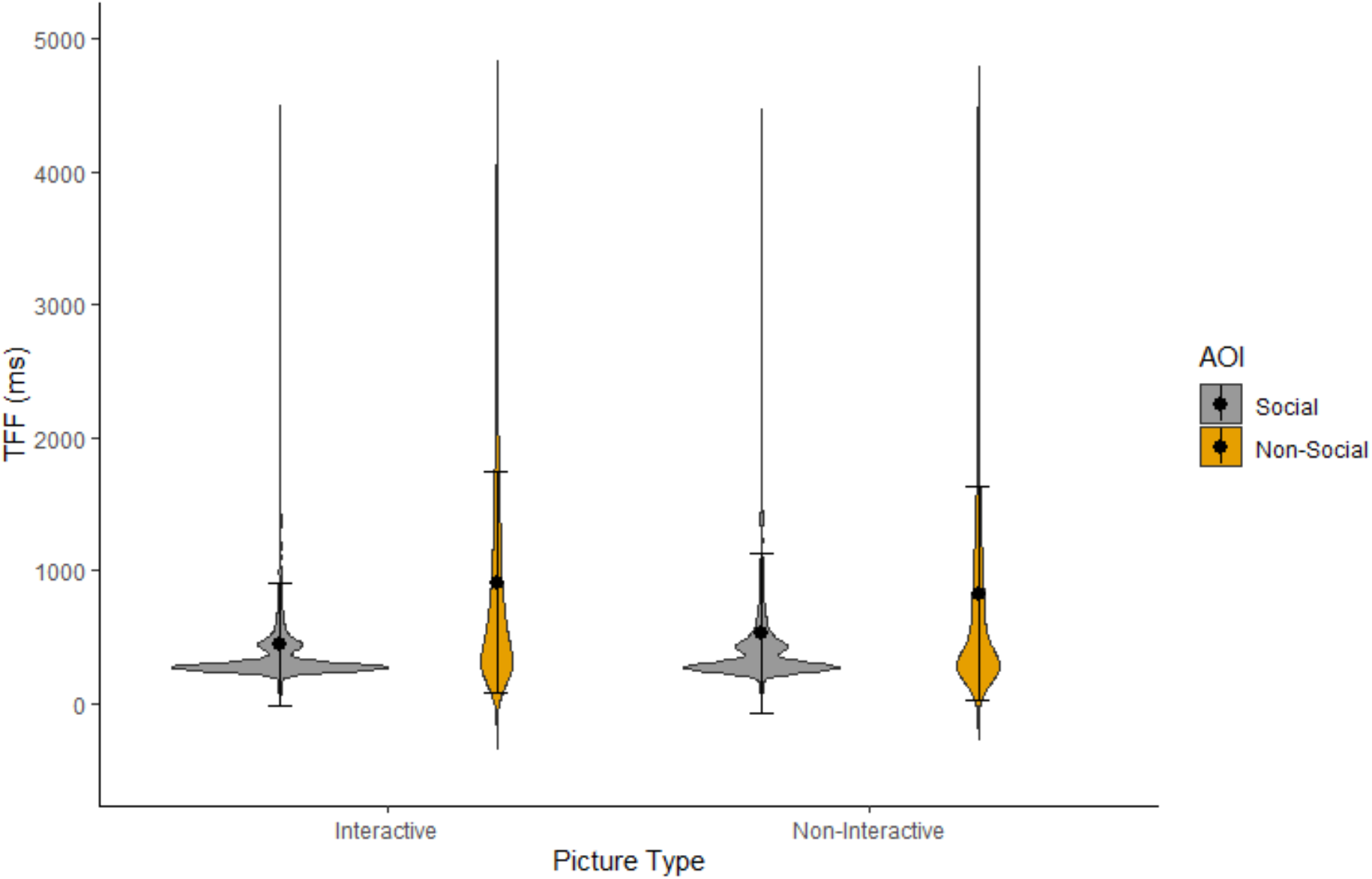
Violin plot for participant’s mean time-to-first-fixation per AOI and scene type. Error bars represent standard deviations.

## Discussion

While much prior research has confirmed the ‘social attention bias’, we demonstrate here that this bias is moderated by the presence (or absence) of social interactions. The currents findings are in line with previous research on both social and non-social attention (e.g. Bindemann, Burton, Hooge, Jenkins, & de Haan, 2005; Bindemann, Scheepers, Ferguson, & Burton, 2010; Fletcher-Watson, Findlay, Leekam, & Benson, 2008; Rigby, Stoesz, & Jakobson, 2016). Indeed, when interactions are not taken into account, our results confirmed previous findings that people orient faster to humans and look at them for longer than other elements in a scene. However, our findings also reveal that this social bias in attentional engagement only occurred when people in the scene were involved in a social exchange. In non-interactive scenes, participants explored the background for just as long as the people. In contrast, participants were faster to orient to human information in *both* interactive and, to a lesser degree, non-interactive scenes.

Most previous research has not directly compared interactive and non-interactive scenes, but instead has looked at attention to isolated individual faces (Bindemann et al., 2005), single person scenes (Bindemann et al., 2010; Fletcher-Watson et al., 2008), or mostly interactive scenes only (e.g. Klin, Jones, Schultz, Volkmar, & Cohen, 2002; Riby & Hancock, 2008; Rigby et al., 2016). Conversely, while we have directly compared noninteractive and interactive scenes, we have not compared attention to dyads to attention to either fewer (i.e. an isolated person) or greater numbers of people. For example, it is possible that an individual may be perceived as more unusual and thus may capture attention more quickly and be processed for longer than two non-interacting individuals; whilst the presence of additional people in a scene may change attention to interactions themselves. The current findings are, however, in line with previous research showing that facing figures are processed differently than non-facing figures (Papeo & Abassi, 2019; Papeo et al., 2019, 2017; Stagg et al., 2014; Vestner, Tipper, Hartley, Over, & Rueschemeyer, 2019). In our view, the current findings further support the notion that the human perceptual system may be at least partially tuned to dyadic social interactions.

Those few studies that have previously explicitly investigated the effect of social interactions on looking behaviour provide inconsistent results (Birmingham et al., 2008; Villani et al., 2015). Our findings are in line with those of Villani et al. (2015), who also found that people looked more at human information in interactive paintings than in noninteractive paintings. Yet, they partially contradict those of Birmingham et al. (2008), who did not observe differences between interactive and individual-action scenes depicting three people. Stimulus and design differences between our study and that of Birmingham et al. (2008) could explain these divergent results. For instance, all our scenes involved two people, while Birmingham et al. (2008) compared scenes with three people to those with one individual. Future research should investigate the possibility that the introduction of additional agents in the scene may lead to changes in attention to and processing of social interactions.

Our findings further our understanding of social scene perception by suggesting that the presence of a social interaction greatly strengthens the social bias as indexed by attentional engagement. We found that the presence of a social interaction fully moderates attentional engagement to social vs. non-social elements of naturalistic scenes. At the same time, this ‘interaction effect’ was not as strong for attentional capture. Although the bias to look first at humans was stronger in scenes that contained a social interaction, participants looked at humans faster than at other elements of the scene whether scenes were interactive or not. Taken together, these findings offer insight into a potential hierarchy of attentional processes that occur when viewing social scenes. Specifically, it lends strength to the idea that the social bias is an automatic bottom-up process that guides our attention, even in cluttered scenes. This is in line with other studies showing faster orienting to human than other information within scenes (Bindemann et al., 2010), quicker fixations on scenes with humans over concurrently presented non-social scenes (Fletcher-Watson et al., 2008), and that social features partially override the impact of low-level physical saliency during scene perception (Rösler, End, & Gamer, 2017). Our results further suggest that the informative value of social interactions guides one’s attention during both ‘first-pass’ rapid orienting and during further exploration once the locations of relevant scene elements have been determined. It is indeed intuitive that we would look at any social information faster than non-social information, but only keep engaged with information of particular social relevance. However, as the current study only compared the existence of a social bias between trials in different conditions, only tentative conclusions can be drawn on prioritization of orienting towards interacting over non-interacting individuals who are present in the same scene. Future studies involving within trial/scene comparisons are necessary in order to further disentangle these processes.

## Author Note

This work has received funding from the European Research Council under the European Union’s Horizon 2020 research and innovation programme (ERC-2016-STG-716974: Becoming Social).

The authors would like to thank Josh Barlow and Clara Hickford Martinez for their help in preparing the stimuli and collecting the data, respectively.

Declarations of interest: none.

## Footnotes

2 Monocular gaze data chosen on the basis of more accurate calibration was used in the analysis.

## References

Bindemann, M., Burton, a. M., Hooge, I. T. C., Jenkins, R., & de Haan, E. H. F. (2005).Faces retain attention. Psychonomic Bulletin & Review, 12(6), 1048–1053. https://doi.org/10.3758/BF03206442

Bindemann, M., Scheepers, C., Ferguson, H. J., & Burton, A. M. (2010). Face, Body, and Center of Gravity Mediate Person Detection in Natural Scenes. Journal of Experimental Psychology: Human Perception and Performance, 36(6), 1477–1485. https://doi.org/10.1037/a0019057

Birmingham, E., Bischof, W. F., & Kingstone, A. (2008). Social attention and real-world scenes: the roles of action, competition and social content. Quarterly Journal of Experimental Psychology (2006), 61(7), 986–998. https://doi.org/10.1080/17470210701410375

End, A., & Gamer, M. (2017). Preferential processing of social features and their interplay with physical saliency in complex naturalistic scenes. Frontiers in Psychology, 8(MAR), 418. https://doi.org/10.3389/fpsyg.2017.00418

Fletcher-Watson, S., Findlay, J. M., Leekam, S. R., & Benson, V. (2008). Rapid detection of person information in a naturalistic scene. Perception, 37(4), 571–583. https://doi.org/10.1068/p5705

Fletcher-Watson, S., Leekam, S. R., Benson, V., Frank, M. C., & Findlay, J. M. (2009). Eyemovements reveal attention to social information in autism spectrum disorder. Neuropsychologia, 47(1), 248–257. https://doi.org/10.1016/j.neuropsychologia.2008.07.016

Kingstone, A. (2009, February 1). Taking a real look at social attention. Current Opinion in Neurobiology. Elsevier Current Trends. https://doi.org/10.1016/j.conb.2009.05.004

Klin, A., Jones, W., Schultz, R., Volkmar, F., & Cohen, D. (2002). Visual Fixation Patterns During Viewing of Naturalistic Social Situations as Predictors of Social Competence in Individuals With Autism. Archives of General Psychiatry, 59(9), 809. https://doi.org/10.1001/archpsyc.59.9.809

Lee, V., & Rutherford, M. D. (2018). Sixteen-month-old infants are sensitive to competence in third-party observational learning. Infant Behavior and Development, 52, 114–120. https://doi.org/10.1016/j.infbeh.2018.07.001

Lenth, R. (2018). Emmeans: estimated marginal means. Aka Least-Squares Means.

Papeo, L., & Abassi, E. (2019). Seeing social events: The visual specialization for dyadic human-human interactions. Journal of Experimental Psychology: Human Perception and Performance, 45(7), 77–888. https://doi.org/10.1037/xhp0000646

Papeo, L., Goupil, N., & Soto-Faraco, S. (2019). Visual Search for People Among People.Psychological Science, 30(10), 1483–1496. https://doi.org/10.1177/0956797619867295

Papeo, L., Nicolas, G., & Hochmann, J.-R. (2020). Visual perception grounding of social cognition in preverbal infants. https://doi.org/10.31219/osf.io/v7y9x

Papeo, L., Stein, T., & Soto-Faraco, S. (2017). The Two-Body Inversion Effect.Psychological Science, 28(3), 369–379. https://doi.org/10.1177/0956797616685769

Pinheiro, J., Bates, D., DebRoy, S., & Sarkar, D. (2016). nlme: Linear and Nonlinear Mixed Effects Models [R package]. R Package Version.

Powell, L. J., & Spelke, E. S. (2018). Human infants’ understanding of social imitation: Inferences of affiliation from third party observations. Cognition, 170, 31–48. https://doi.org/10.1016/j.cognition.2017.09.007

Quadflieg, S., & Koldewyn, K. (2017). The neuroscience of people watching: How the human brain makes sense of other people’s encounters. Annals of the New York Academy of Sciences. https://doi.org/10.1111/nyas.13331

Quadflieg, S., & Penton-Voak, I. S. (2017). The Emerging Science of People-Watching:Forming Impressions From Third-Party Encounters. Current Directions in Psychological Science. https://doi.org/10.1177/0963721417694353

Riby, D. M., & Hancock, P. J. B. (2008). Viewing it differently: social scene perception in Williams syndrome and autism. Neuropsychologia, 46(11), 2855–2860. https://doi.org/10.1016/j.neuropsychologia.2008.05.003

Rigby, S. N., Stoesz, B. M., & Jakobson, L. S. (2016). Gaze patterns during scene processing in typical adults and adults with autism spectrum disorders. Research in Autism Spectrum Disorders, 25, 24–36. https://doi.org/10.1016/j.rasd.2016.01.012

Risko, E. F., Laidlaw, K. E. W., Freeth, M., Foulsham, T., & Kingstone, A. (2012, May 25). Social attention with real versus reel stimuli: Toward an empirical approach to concerns about ecological validity. Frontiers in Human Neuroscience. Frontiers Media S. A. https://doi.org/10.3389/fnhum.2012.00143

Rösler, L., End, A., & Gamer, M. (2017). Orienting towards social features in naturalistic scenes is reflexive. PLoS ONE, 12(7), e0182037. https://doi.org/10.1371/journal.pone.0182037

SR Research Ltd. (2012). Eyelink Programmers Guide.

SR Research Ltd. (2013). EyeLink Data Viewer User’s Manual (No. 1.11.900). Mississauga, Canada.

Stagg, S. D., Linnell, K. J., & Heaton, P. (2014). Investigating eye movement patterns, language, and social ability in children with autism spectrum disorder. Development and Psychopathology, 26(02), 529–537. https://doi.org/10.1017/S0954579414000108

Vestner, T., Tipper, S. P., Hartley, T., Over, H., & Rueschemeyer, S. A. (2019). Bound together: Social binding leads to faster processing, spatial distortion, and enhanced memory of interacting partners. Journal of Experimental Psychology: General, 148(7), 1251–1268. https://doi.org/10.1037/xge0000545

Villani, D., Morganti, F., Cipresso, P., Ruggi, S., Riva, G., & Gilli, G. (2015). Visual exploration patterns of human figures in action: An eye tracker study with art paintings. Frontiers in Psychology, 6(OCT), 1636. https://doi.org/10.3389/fpsyg.2015.01636

Xiao, J., Hays, J., Ehinger, K. A., Oliva, A., & Torralba, A. (2010). SUN database: Large-scale scene recognition from abbey to zoo. In 2010 IEEE Computer Society Conference on Computer Vision and Pattern Recognition (Vol. 29, pp. 3485–3492). IEEE. https://doi.org/10.1109/CVPR.2010.5539970

Yarbus, A. (1967). Eye movements and vision. New York, NY: Plenum Press.

